# Phosphorylation by the PknB serine/threonine kinase stimulates dimer formation by WalR

**DOI:** 10.1101/2025.07.12.664526

**Authors:** Marina Suppi, Ian R. Monk, Liam K. R. Sharkey, Nichollas E. Scott, Dweipayan Goswami, Aakash Natarajan, Stephanie Tan, Katharine Myler, Sheila Marie Pimentel-Elardo, Timothy P. Stinear, Sacha J. Pidot, Justin R. Nodwell

## Abstract

WalKR is a two-component system that regulates cell wall homeostasis and other processes in low-GC Gram-positive bacteria. It is essential for viability in *Staphylococcus aureus* and regulates genes that encode the autolysins and other critical proteins. The sensor kinase WalK phosphorylates WalR at aspartic acid residue 53 (D53) in the receiver domain, stimulating promoter DNA binding. Unlike other response regulators, WalR is thought to have a second kinase, the serine/threonine kinase PknB, which phosphorylates the receiver domain at threonine residue 101. We previously reported that a *walR* mutation that changed T101 to a methionine conferred low level resistance to vancomycin and greatly increased susceptibility to tunicamycin along with several other phenotypic traits. In this work we demonstrate similarities between *pknB* null and the WalR_T101M_ mutants that support a regulatory role for PknB-mediated phosphorylation of WalR. Using a combination of *in vitro* and *in vivo* approaches, we confirm the specificity of this posttranslational modification and confirm that phosphorylation at both D53 and T101 is important for dimer formation. Importantly, using an *in silico* approach, we have discovered that phosphorylation at T101 creates an intermolecular hydrogen bond between the phosphate group and E108, that strengthens contacts along the WalR dimerization interface. This is the first time that a clear molecular role has been assigned to this posttranslational modification and it is the strongest molecular evidence to date for a direct functional relationship between PknB and WalR. We propose that, together with primary activation by WalK, PknB serves as a second potentiating input into the WalKR system, stimulating WalR dimer formation, and target DNA binding and tethering this transcriptional event to important extracellular stimuli.

## Introduction

Antimicrobial resistance in pathogenic bacteria, threatens human health and healthcare systems worldwide (1,2). One of the most common causes of post-operative infection, *Staphylococcus aureus,* has emerged in resistant forms including <u>m</u>ethicillin-resistant *<u>S</u>taphylococcus <u>a</u>ureus* (MRSA), <u>v</u>ancomycin intermediate *<u>S</u>taphylococcus <u>a</u>ureus* (VISA) and <u>v</u>ancomycin-resistant *<u>S</u>. <u>a</u>ureus* (VRSA) (1,3). VISA strains have been linked with chronic infection, hospitalization, prolonged vancomycin treatment and/or treatment failure though they are not as aggressive a clinical burden as MRSA or VRSA (1,3). However, VISA-like phenotypes have also appeared in combination with dalbavancin and methicillin resistance, giving rise to strains that are harder still to treat (3,4). Understanding the genetic factors that contribute to these diverse antibiotic-related phenotypes is very important as the resulting insights will reveal new therapeutic strategies (5).

There are several molecular mechanisms that can confer the VISA phenotype (6–10). One determinant is the WalKR two-component system (9,11–13), also referred to as YycFG and VicRK. WalKR is highly conserved in low G+C Gram-positive bacteria and consists of the sensor histidine kinase WalK and the OmpR/PhoP family response regulator WalR. WalKR regulates genes involved in cell wall homeostasis (14–16) including the autolysins (*atl*, *sle1*, *sceD*, *lytM*, *isaA* and *ssaA*) which cleave bonds in the peptidoglycan permitting cell growth and division (17). It also regulates *hup* and *dnaA,* involved in DNA compaction and replication (18) and genes involved in lipoteichoic acid synthesis (*ltaS*), translation (*rplK*) and purine nucleotide metabolism (*prs*) (18). The direct regulation of autolysins and other cell wall-acting genes is consistent with the involvement of *walKR* mutations in intermediate resistance to vancomycin, a thickened cell wall, reduced autolytic activity and reduced virulence, all of which are hallmarks of VISA (9,11–13).

Like most histidine kinases, WalK autophosphorylates at a conserved histidine residue and then transfers that phosphate group to a conserved aspartate in the WalR receiver domain, stimulating DNA binding at target promoters (19,20). Unlike the other 15 two-component systems in *S. aureus,* WalKR is essential for viability (11,21), consistent with its central physiological role. The most unusual feature of this system, discovered by the Dworkin and Grein groups, is that the serine/threonine kinase PknB (PrkC in *Bacillus subtilis*) phosphorylates WalR at residue T101 (22,23). T101 is in the “α4-β5-α5” moiety of the receiver domain which makes up most of the protein’s dimerization interface. Response regulators having multiple histidine kinases are known (24), however, mostly they are histidine kinases that phosphorylate the conserved regulatory aspartate in the receiver domain. In contrast, the phosphorylation of WalR by a ser/thr kinase at the T101 residue is unique to this family.

Like many response regulators, WalR consists of an N-terminal receiver domain, which includes the conserved WalK phosphorylation target at residue D53, and a C-terminal DNA binding domain. Structural and biochemical studies have suggested that phosphorylation at the conserved receiver domain aspartate modulates dimerization in many other response regulators. For example, *S. aureus* protein VraR is monomeric in the unphosphorylated form and occupies a ‘closed’ conformation that cannot bind DNA. In this conformation the DNA binding domain occludes α4 and β5 in the α4-β5-α5 dimerization interface. Phosphorylation by its cognate histidine kinase triggers the open conformation, permitting dimerization and, thereby, the capacity to bind target promoter DNA (25). A similar model has been proposed for the *E. coli* OmpR protein and other well-characterized response regulators (26–28). There are several structural variations on this theme, but dimer formation is clearly a central regulatory feature of response regulators (29). WalR is not understood at this level of detail, however there is evidence that phosphorylation at D53 stimulates dimerization and DNA binding (18,30). In contrast, the role of phosphorylation at T101 is unknown.

In previous work, we characterized a number of point mutations in *walKR* that confer resistance to the cell wall-acting antibiotics actinorhodin and siamycin I (31,32), using the so-called “Rosenbach” type strain, ATCC29213. One of these mutations, which we referred to as *walR1,* was of particular interest because it changed T101 to a methionine. We will refer to this mutation as *walR_T101M_* in this work so that we can more clearly discuss it in the context of other alleles. We characterized *walR_T101M_* in detail and showed that it conferred a VISA-like phenotype. This included low level resistance to vancomycin and ramoplanin, a thickened cell wall and a significant defect in autolysis. A striking feature of this mutation is that it also conferred a pronounced sensitivity to tunicamycin. Tunicamycin is a substrate mimetic of the enzymes MraY and TarO, which catalyze the first steps in peptidoglycan and teichoic acid biosynthesis, respectively (16,33–35). We were able to attribute this to the partial collapse of a peptidoglycan recycling pathway carried out by the autolysins and the products of the *mupG* peptidoglycan recycling operon (16). A logical next step in this work was to investigate this mutation, and others at T101, in greater detail to determine their molecular effect on WalR function. The key question being whether the VISA-like phenotype was due to a failure in phosphorylation by PknB, a structural alteration resulting from the T101M replacement, or both.

PknB is composed of three extracellular PASTA domains (for <u>P</u>enicillin-binding protein <u>a</u>nd <u>S</u>erine/<u>T</u>hreonine kinase <u>A</u>ssociated domain), a central transmembrane domain and an intracellular kinase domain (36). PknB phosphorylates WalR via direct transfer of the γ-phosphate of ATP to a threonine (T101) in the receiver domain (37). PknB regulates critical cellular processes in *S. aureus*, including cell wall metabolism and division (16,18,23,38), antibiotic resistance (16,18,38–40), purine biosynthesis (38), virulence (38,40,41), and central metabolism (38). Notably, a deletion of *pknB* leads to increase sensitivity to cell wall-active antibiotics such as β-lactams or tunicamycin (38,40). PknB localizes to the septum of the cell, and it has been reported to phosphorylate the early cell division protein FtsZ (23). The kinase also regulates purine biosynthesis through direct phosphorylation of adenylosuccinate synthase (PurA), and serves as a positive regulator of the alternative sigma factor SigB, a transcription factor responsible for promoting expression of genes that confer resistance to stress responses (40,42). Its link to the cell wall is further demonstrated by the presence of the three extracellular PASTA domains which bind to muropeptides, components of the cell wall, and lipid II, a precursor for peptidoglycan synthesis, in *M. tuberculosis* and *S. aureus* (23,43,44). Work in *Bacillus subtilis* (22), showed that PrkC, the PknB homolog, phosphorylated WalR in *in vitro* assays and that this occurred at T101. This was corroborated *in vivo* using phos-tag analysis and mass spectrometry. It was also reported that a *pkrC* deletion altered the expression of some WalR target genes, including *iseA* and *pdaC* (22). Similar work in *S. aureus* showed that PknB phosphorylates WalR *in vitro* at T101 (23), and the deletion of *pknB* causes a change in the expression of WalR target genes, including some autolysins (38). Thus, the biological roles attributed to PknB are consistent with involvement in the WalKR system however, again, the biochemical role of this modification is unknown.

In this work, we explore the role of PknB and the T101 residue in WalR in *S. aureus* ATCC29213. We find that deletion of *pknB* alters several cellular features including alterations in antibiotic sensitivity and autolytic activity. These changes are consistent with a role in the modulation of signaling through WalKR. We further show that phosphorylation of WalR at T101 promotes dimer formation *in vitro* and *in vivo*. *In silico* modelling and *in vivo* assays suggest that stabilization of the dimer is mediated by an intermolecular hydrogen bond between phosphorylated T101 and E108, another residue in the α5-β4-α5 region of the receiver domain. We suggest that this mechanism of dimer stabilization provides a secondary regulatory input into this important system.

## Results and Discussion

### A complex relationship between the Δ*pknB* and WalR_T101M_ phenotypes

The *walR*_T101M_ point mutation confers a VISA-like phenotype in *S. aureus* ATCC29213 including 2-4x increased MIC for vancomycin, a thickened cell wall, and a defect in autolysis (16). This mutation also confers a 64x increase in sensitivity to the antibiotic tunicamycin (16). This “collateral sensitivity” is unusual and merits further investigation as it could constitute an Achilles heel in an otherwise antibiotic-resistant pathogen. To initiate this investigation, we constructed a *pknB* null mutation (Δ*pknB*) in *S. aureus* ATCC29213 to determine whether it would phenocopy or partially phenocopy the effects of *walR_T101M_*.

We determined the MICs of antibiotics that act on the five major antibiotic targets, comparing Δ*pknB* to its wild type parent (**Table 1**). Deletion of *pknB* had no detectable effect on sensitivity to most drugs that target the cell wall (including vancomycin and fosfomycin), DNA gyrase (ciprofloxacin), RNA polymerase (rifampicin), and the ribosome (kanamycin). However, *pknB* deletion conferred a two-fold increase in sensitivity to oxacillin and an eight-fold increase in sensitivity to tunicamycin. These features are shared with WalR_T101M_ though the tunicamycin sensitization is less pronounced (8-fold vs 64-fold; **Table 1**). Complementation of the Δ*pknB* mutation *in trans* restored normal sensitivity to tunicamycin, demonstrating that the phenotype was due to the *pknB* deletion.

**Table 1.**
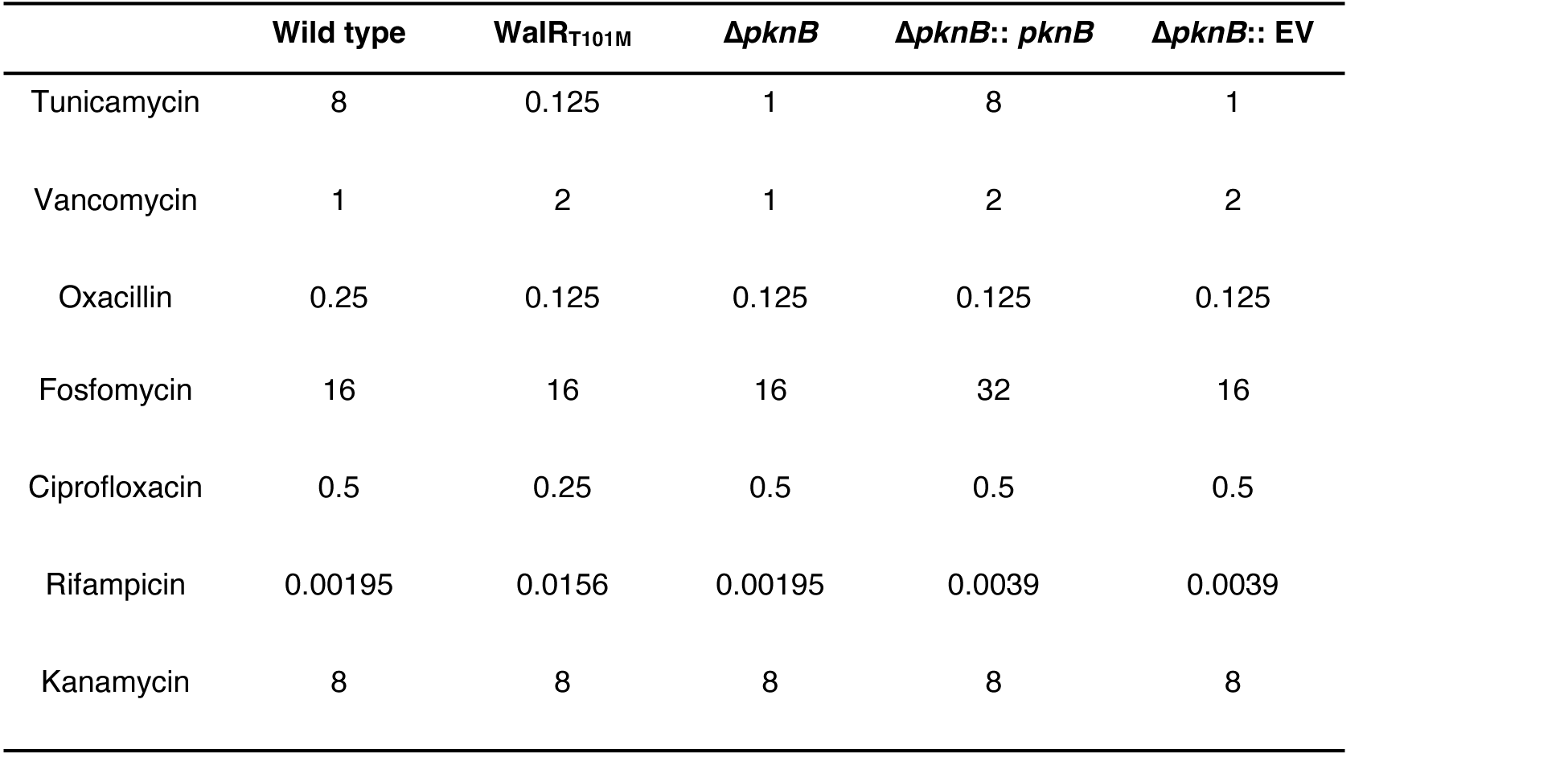
MICs of each strain against a panel of antibiotics. Drug concentrations are expressed in μg/mL. The results are a representation of three independent biological replicates.

| | Wild type | WalR<sub>T101M</sub> | Δ*pknB* | Δ*pknB*:: *pknB* | Δ*pknB*:: EV |
| --- | --- | --- | --- | --- | --- |
| Tunicamycin | 8 | 0.125 | 1 | 8 | 1 |
| Vancomycin | 1 | 2 | 1 | 2 | 2 |
| Oxacillin | 0.25 | 0.125 | 0.125 | 0.125 | 0.125 |
| Fosfomycin | 16 | 16 | 16 | 32 | 16 |
| Ciprofloxacin | 0.5 | 0.25 | 0.5 | 0.5 | 0.5 |
| Rifampicin | 0.00195 | 0.0156 | 0.00195 | 0.0039 | 0.0039 |
| Kanamycin | 8 | 8 | 8 | 8 | 8 |

In addition to the altered sensitivity to cell wall-active antibiotics, the *walR_T101M_* allele confers significant changes in cell wall homeostasis (11,14,16,18) including increased peptidoglycan thickness (16). When examined under transmission electron microscopy, we observed no significant difference in thickness of the Δ*pknB* cell wall (30.7 ± 5.3 nm compared to 35.4 ± 7.4 nm in the wild type) (**Figure 1B** and **Figure S1**) (12,16). We also saw no difference in cell division status of Δ*pknB* compared to the parent strain in either lag or stationary phase (**Figure 1C**). Thus, Δ*pknB* and WalR_T101M_ differ considerably in cell morphology.

**Figure 1.**
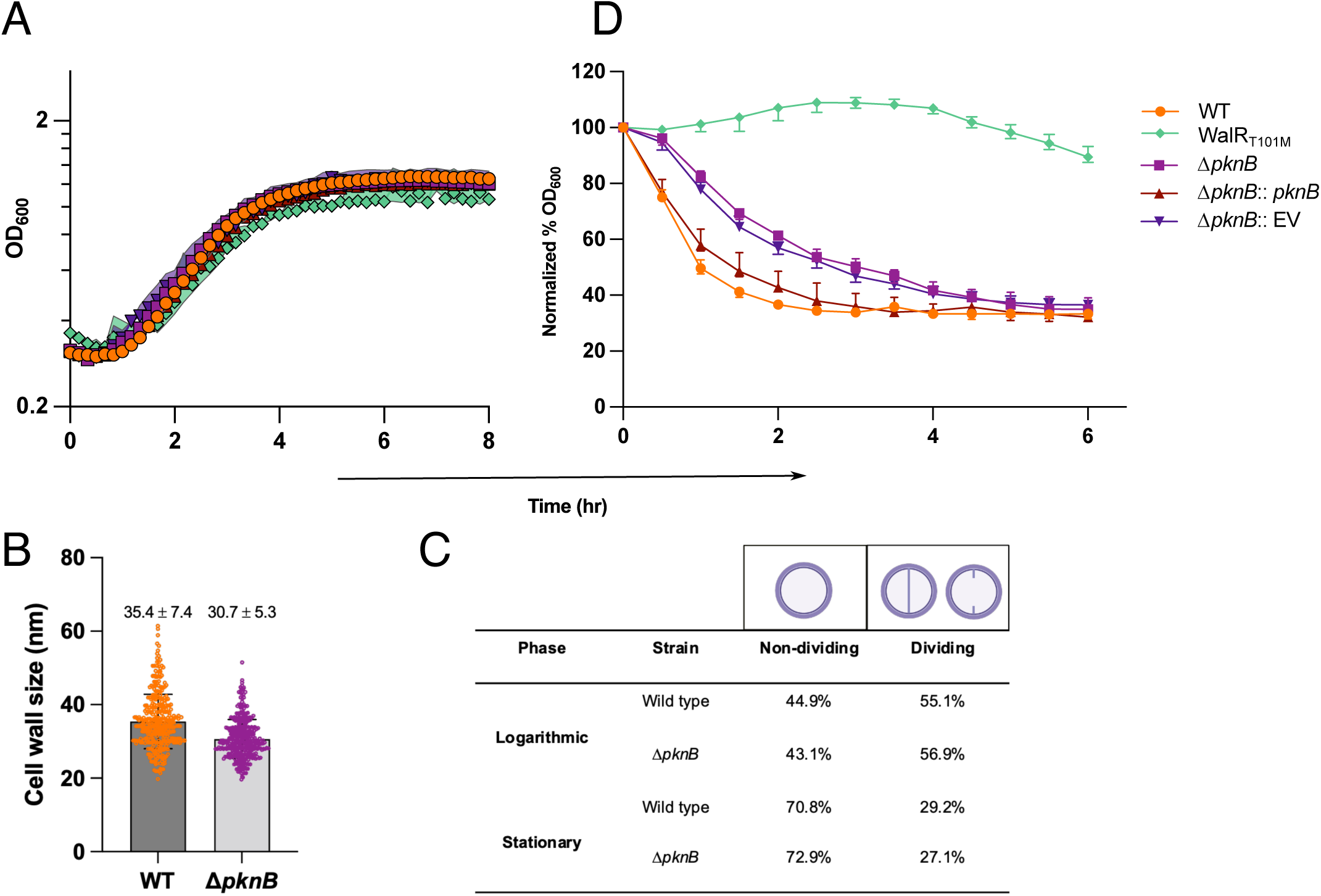
Phenotypic study of the Δ*pknB* mutant. **A)** Growth curves expressed in OD600 nm over a period of 8h. **B)** Cell wall thickness of wild type and **Δ***pknB* strains assessed by measuring over 350 cells. **C)** Percentage of cells in either non-dividing or dividing state based on the 350 cells from transmission electron microscopy. **D)** *In vivo* autolysis of WalKR mutant strains. Cells were exposed to 0.05% (v/v) Triton X-100 detergent and autolytic activity was measured by change in turbidity at OD600 nm over a period of 6h.

WalKR directly regulates the autolysin-encoding genes (14,15,18) so we compared the autolytic activity of each strain (**Figure 1D**). Wild type cells displayed rapid autolysis; culture turbidity decreased by 71.6% over a 6h-period. Consistent with our previous results, the WalR_T101M_ mutant was defective in autolysis, with culture turbidity decreasing by only 10.6% over 6 h. The Δ*pknB* mutant exhibited a reproducible delay in autolysis, but reached the same level of cell lysis as wild type after 6 hrs, consistent with existing data in other strains (38). As above, complementation of Δ*pknB in trans* restored autolysis to the wild type profile (**Figure 1D**).

In sum, the *pknB* deletion did not phenocopy *walR_T101M_*mutation, although their phenotypic traits exhibited considerable overlap. A possible explanation could be that the WalR_T101M_ mutant suffers both from lack of phosphorylation by PknB and from a subtle structural defect caused by the amino acid substitution. Protein folding and stability would need to be sufficient to permit baseline function, including phosphorylation by its cognate histidine WalK, since both *walK* and *walR* are essential. This hypothesis is consistent with the shared sensitivity to tunicamycin and defect in autolysis, and with the differing severity in the two mutants.

### Phosphorylation at T101 stimulates WalR dimer formation

The phenotypic comparison of the Δ*pknB* and WalR_T101M_ mutants was consistent with a regulatory role for PknB-mediated phosphorylation of WalR. To address this relationship more directly, we used biochemical and two-hybrid approaches to directly investigate the effect of PknB-mediated phosphorylation on WalR.

We expressed and purified the cytoplasmic domain of PknB (residues 1-348, PknB^CYTO^) along with WalR including the wild type, T101M, T101A, and T101S mutants. We carried out *in vitro* time course γ-^32^P-ATP phosphorylation assays. Incubation of PknB^CYTO^ with wild type WalR resulted in phosphorylation of WalR within 5 min. Consistent with previous work suggesting that T101 is the target for this phosphorylation, we found that all amino acid substitutions at this residue eliminated this modification (**Figure 2**). Ser/thr kinases can phosphorylate both threonine and serine residues, thus it is particularly striking that even the T101S mutation blocked this modification. This suggests that phosphorylation by PknB is highly specific for the threonine residue (**Figure 2**).

**Figure 2.**
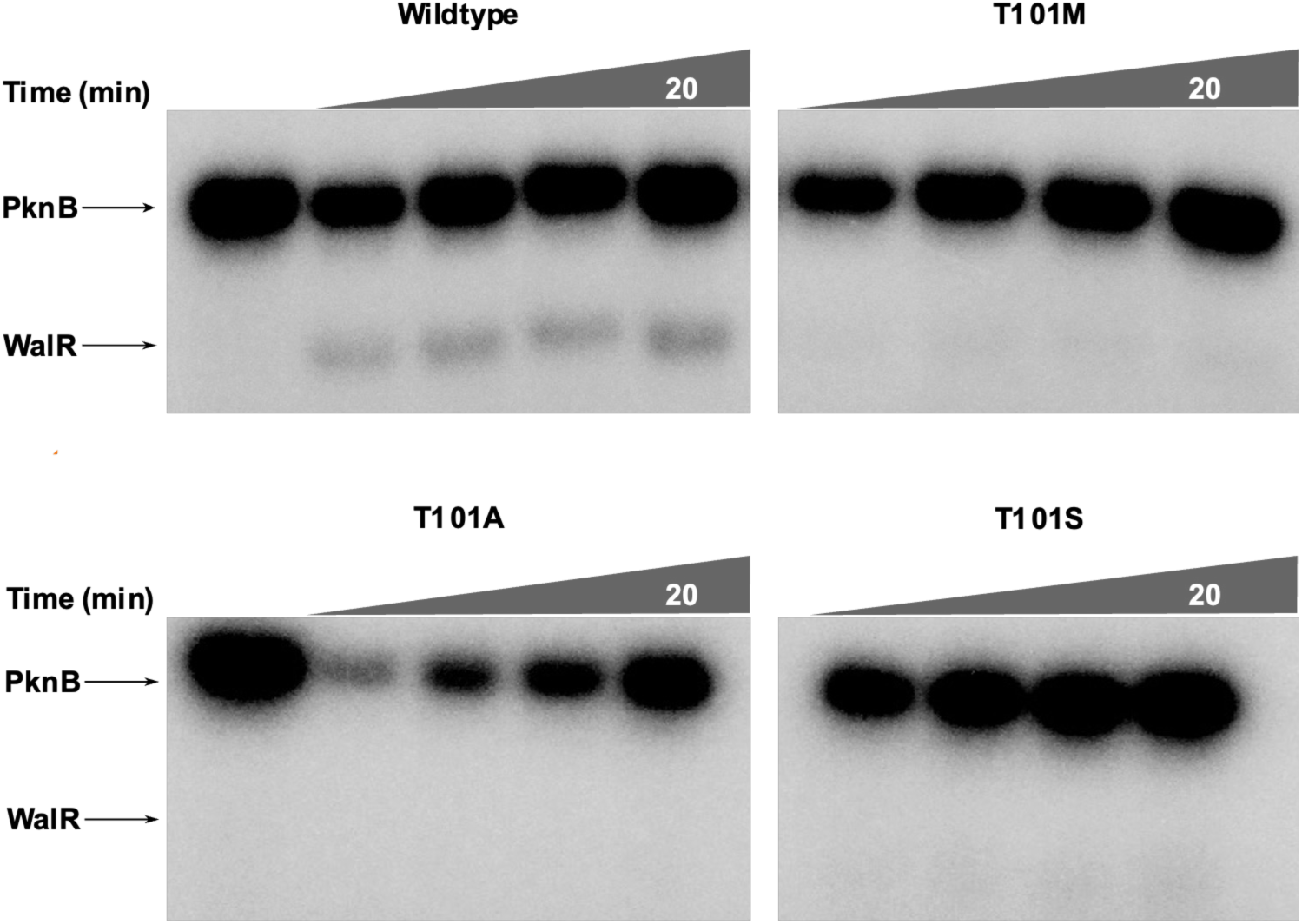
*In vitro* phosphorylation assays with four variants of WalR and PknB. Full-length wild type WalR and three other variants (WalRT101M, WalRT101A and WalRT101S) were incubated with PknB (cytoplasmic part only) in phosphorylation buffer in the presence of 2.5 μCi [ψ-32P] ATP for 20 minutes (1:4 ratio; kinase: regulator). Samples were analyzed in a 12% SDS-PAGE gel, followed by autoradiography. Autoradiographs shown are a representation of three independent experiments.

WalR is a conventional response regulator of the OmpR/PhoP subfamily with a winged helix-turn-helix DNA-binding domain (29,45). It forms a head-to-head dimer of the receiver domains, using the conserved α4-β5-α5 face for intermolecular interactions (**Figure 4A**), and a head-to-tail dimer of the winged helix-turn-helix motifs that bind to the tandem DNA repeats of its binding site (46). As the T101 residue falls in the dimerization interface, we asked whether this residue, and/or its phosphorylation might influence dimer formation. To address this hypothesis, we used size-exclusion chromatography multi-angle light-scattering (SEC-MALS) to assess the monomer/dimer status of wild type and mutant WalR proteins.

We compared the oligomeric status of WalR in the presence of PknB^CYTO^, with or without ATP. In the absence of ATP, WalR was predominantly monomeric. Addition of ATP shifted WalR into a predominantly dimeric form. When we repeated this experiment with WalR_T101M_ we found that the protein behaved as a monomer, regardless of the addition of ATP (**Figure 3A** and **Table S3**). The T101M mutation did not influence the thermal melt properties of the protein, indicating that any structural reorganization caused by this mutation must be quite subtle (**Figure S2**). Aside from this, this result is consistent with the idea that phosphorylation of T101 stabilizes the WalR dimer.

**Figure 3.**
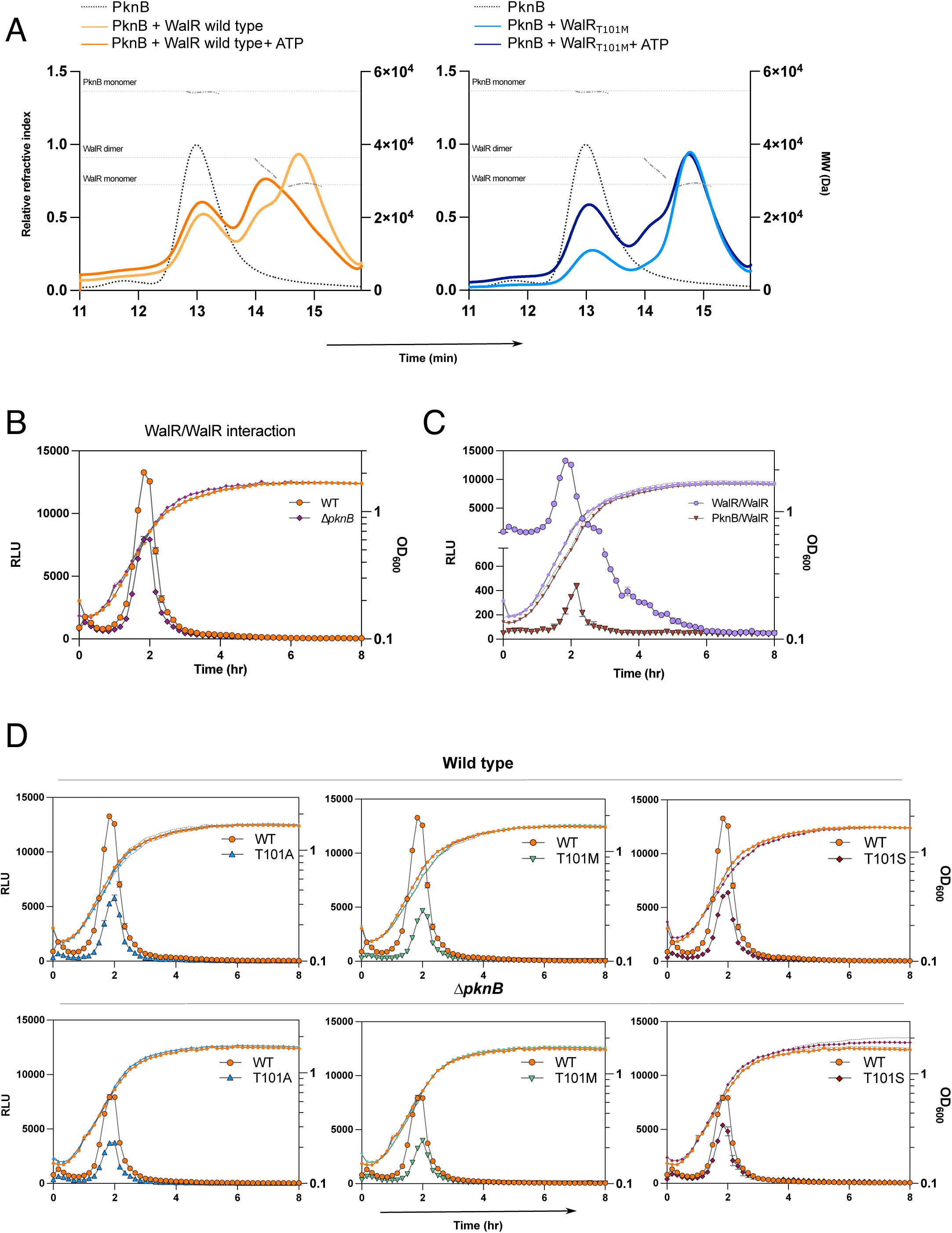
Dimerization studies and kinetic interactions of wild type and WalR mutants in *S. aureus.* **A)** Elution profiles of wild type WalR and WalRT101M proteins upon phosphorylation by PknB analyzed by SEC-MALS. Samples are labelled as follows: orange lines are for wild type protein, and blue are for WalRT101M; light orange or blue how reaction without ATP, and dark orange or blue show with the addition of ATP. Elution profile of PknB alone is depicted by a grey dashed line. The dotted line at each peak indicates the measured molecular weight (MW). The dotted horizontal grey lines along the right *y*-axis represent the theoretical MW of the monomer and dimer of he indicated proteins. Results are the median from three independent assays. **B)** Analysis of the interaction of WalR-SmBIT/LgBIT throughout growth in wild type and Δ*_pknB_* backgrounds. **C)** Kinetic interactions of WalR/WalR and WalR/PknB throughout growth in wild type cells. **D)** Comparison of wild type WalR/WalR kinetics to three mutant versions of WalR (WalRT101M, WalRT101A, and WalRT101S) in wild type and Δ*_pknB_* backgrounds. **B-D)** Results represent one biological replicate and error bars or dashed lines show the error range (minimum to maximum values). Two other biological replicates are shown in the Supplemental material **Figure S4**.

As a second approach, we took advantage of a split luciferase assay developed for *S. aureus* (18). To directly probe WalR dimerization *in vivo,* we constructed WalR fusions to the two fragments of the Nanoluc® luciferase “SmBIT” and “LgBIT” and introduced them into the wild type and the Δ*pknB* mutant. In the wild type background, we observed a peak luciferase signal in the early-exponential phase of growth, indicating complex formation by the two fusion proteins resulting from WalR dimerization. The signal subsequently declined to undetectable levels during stationary phase (**Figure 3B**). We observed a similar peak in the Δ*pknB* mutant however the signal was reduced by 36.5% compared to the wild type (**Figure 3B**), suggesting that PknB is important but not essential for dimer formation by WalR. It was not possible to assess this interaction in a *walK* null mutant because the kinase is essential for viability. Nevertheless, we surmise that the remaining dimer formation was due to phosphorylation by the histidine kinase at D53.

Subsequently, we probed the PknB/WalR interaction using a PknB-LgBIT fusion in combination with WalR-SmBIT. In this experiment we observed a peak of PknB/WalR interaction that occurred simultaneously with the WalR/WalR dimerization peak, in early-exponential phase (2h) (**Figure 3C**). This is consistent with the idea that PknB phosphorylates WalR at T101 and that this modification is coincident with WalR dimer formation. We then introduced T101M, T101A and T101S mutations and examined their effect on luciferase readouts. The T101M mutation caused a 65.2% decrease in peak luciferase signal area in the WalR-Smbit/WalR-Lgbit strain. The T101A and T101S mutations also reduced light emission, though to a lesser extent, with a 56.7% and 52.4% decrease in peak area compared to wild type proteins, respectively (**Figure 3D**). These results are consistent with a requirement for T101 for a stable *in vivo* interaction between PknB and WalR. They are also consistent with the hypothesis that the T101M mutation has an additional structural influence on WalR function over and above its defect in PknB-mediated phosphorylation.

As an alternative approach, we repeated these experiments in the Δ*pknB* background. We observed comparable peak luciferase signals of the T101M mutant alleles in both the Δ*pknB* and wild type backgrounds (3654 and 4336 RLU, respectively). The same was observed for the T101A and T101S mutations. The only notable difference between the two strains was a signal reduction in the wild type WalR/WalR interaction in the Δ*pknB* background, which we mention above (**Figure 3D**). The interaction profiles of the T101M, T101A, and T101S mutants are similar in both the wild type and Δ*pknB* backgrounds, consistent with the fact that PknB cannot phosphorylate these mutant proteins. In contrast, phosphorylation occurs when threonine is present, which explains why only the wild type interaction is affected by the absence of PknB. Pairing the wild type WalR protein with the T101M allele did not rescue this defect (WalR_T101M_-SmBIT and WalR_WT_-LgBIT) and, similarly, resulted in a 56.5% decrease in peak area compared to the wild type interaction (**Figure S3**).

Together, these observations support two key conclusions: the phosphorylation of WalR at T101 residue by PknB promotes dimerization *in vitro* and *in vivo*; and the methionine replacement at T101 disrupts phosphorylation and also impairs dimer formation by the unphosphorylated protein. These observations support our hypothesis that the WalR_T101M_ phenotype reflects the loss of PknB phosphorylation and a subtle structural alteration that affects dimerization. Most importantly, it is consistent with a model in which phosphorylation of T101 by PknB stimulates the formation of a WalR dimer thereby influencing the expression of WalR target genes.

### Phosphorylated T101 creates an intermolecular hydrogen bond with E108

To further understand the role of T101 in WalR dimerization, we generated an *in silico* structural model of the dimerization interface between WalR monomers. The amino acid sequence of the WalR receiver domain (residues 1-122), including the dimerization interface, from *S. aureus* strain N315 (UniProtKB entry Q7A8E1) was used to generate a structural model with Swiss-Model web server (47). Notably, WalR from *S. aureus* strain N315 is identical to WalR from *S. aureus* ATCC29213. The resolved crystal structure of WalR receiver domain from *Bacillus subtilis* (PDB: 2ZWM), which is also dimerized, was used as template for accurate modelling (**Figure 4A**). The quality and reliability of the *in silico* WalR receiver domain dimer model were supported by Ramachandran plot analysis, MolProbity, and QMEANDisCo scoring, all of which exceeded accepted thresholds for high-quality structural models (**Figure S5**).

**Figure 4.**
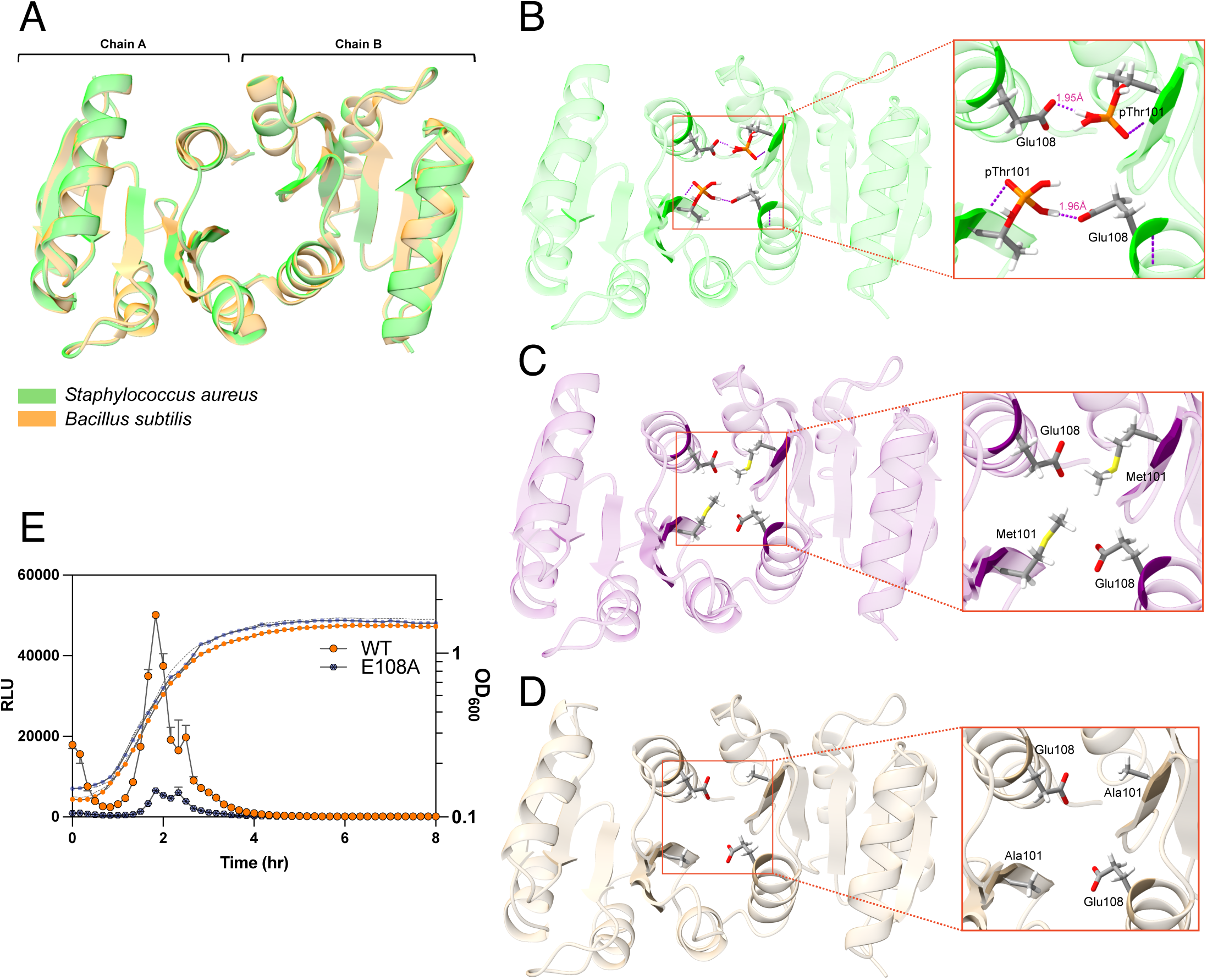
Molecular modelling of WalR receiver domain and T101 interactions in *S. aureus*. **A)** Structural model of a dimer of WalR receiver domain of *S. aureus* superposed with the resolved WalR receiver domain structure of *B. subtilis* (PDB: 2ZWM). *S. aureus* prediction is indicated in green and the resolved structure from *B. subtilis* is shown in orange. Chain A and B represent each a monomer of WalR receiver domain. **B)** Inter-chain interaction analysis of wild type phosphorylated WalR (green), WalRT101M (purple) and WalRT101A (beige) proteins. Phosphorylated T101 residue or its variants are shown as sticks, with carbon atoms illustrated in grey, oxygen in red, hydrogen in white, phosphorus in orange, and sulphur in yellow. Hydrogen bonds are shown as purple dashed lines, and the bond length is indicated in pink. **E)** Comparison of wild type WalR/WalR kinetics to WalRE108A mutant in **Δ***pknB* background. Results represent one biological replicate and error bars, or dashed lines show the error range (minimum to maximum values). Two other biological replicates are shown in the Supplemental material **Figure S6**.

To investigate the role of T101 we performed interaction analysis, focusing on the phosphorylated T101 residue and the α4-β5-α5 dimerization interface. In the model, the phosphate group on T101 formed an intermolecular hydrogen bond with E108 of the neighboring chain, with a bond length of 1.96 Å, well within the expected range (**Figure 4B**). The close fit between the two chains, as shown by solvent-accessible surface area (SASA) analysis, suggests that they pack tightly together (**Supplementary Data T1**). Substituting for a methionine residue (WalR_T101M_) eliminated the hydrogen bond with E108. In turn, the E108 instead formed a hydrogen bond with tyrosine 99 (bond length of 3.1 Å) and a salt bridge to lysine 88 (**Supplementary Data T1**). The absence of polar interactions reduced the buried SASA score at the interface, suggesting decreased interfacial stability (**Figure 4C**). The same could be observed when T101 was substituted for an alanine (WalR_T101A_). Alanine’s small and non-polar nature eliminated hydrogen bonds and salt bridges at residue 101 (**Figure 4D**).

We used our split-luciferase reporter assay to investigate this putative interaction. We generated fusions of WalR_E108A_ to Sm-Bit and Lg-Bit and introduced them into *S. aureus.* Analysis of luciferase signal from co-expression of these two WalR proteins showed a dramatic, 83.7% drop in signal intensity between WalR_E108A_ monomers compared to wild type WalR (**Figure 4E**), suggesting that E108 is important for dimer formation by WalR. This supports our *in silico* model and suggests that the T101∼P-E108 interaction is important for dimer formation between WalR monomers.

To compare the importance of PknB-mediated phosphorylation at T101 with WalK-mediated phosphorylation at D53, we introduced a D53A mutation in our split-luciferase system. The D53A mutation conferred a dramatic reduction in dimer formation in comparison to the wild type interaction. Notably, this reduction was of a similar magnitude to that caused by the T101A mutation (**Figure 5A**). We created double mutants that eliminated both phosphorylation sites, WalR_D53A/T101A_. In this case, signal intensity from the split luciferase system dropped, essentially, to background with a 91.6% decrease in peak signal area compared to wild type interaction (**Figure 5A**). Finally, we examined the effects of the D53A and T101A mutations on dimer formation in the Δ*pknB* background. In this case, we found, again, that only the interaction between the wild type proteins was seriously compromised compared to the wild type strain, supporting a role for phosphorylation at T101 in WalR dimer formation (**Figure 5B**).

**Figure 5.**
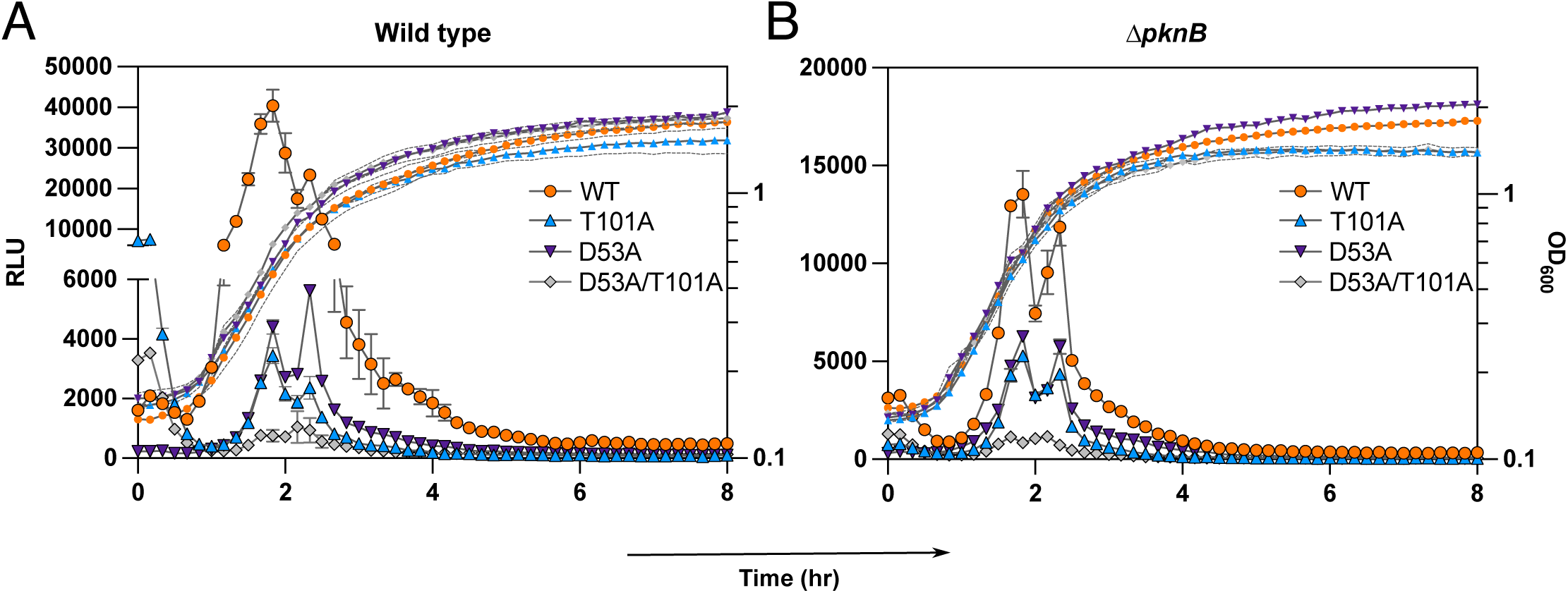
Effects of PknB and/or WalK phosphorylation to WalR dimer formation in *S. aureus.* **A and B)** Comparison of wild type WalR/WalR kinetics to three mutant ersions of WalR (WalRT101A, WalRD53A, and WalRD53A/T101A) in wild type and **Δ***pknB* backgrounds. Results are the representation from three independent assays and error ars, or dashed lines show the error range (minimum to maximum values). Two other biological replicates are shown in the Supplemental material **Figure S7**.

This is the strongest molecular evidence to date for a direct, functional relationship between PknB and WalR. We note that, unlike *walK, pknB* is dispensable for viability in *S. aureus.* We suggest therefore that the critical modification is WalK-mediated phosphorylation of D53 and that, for this reason, cells are viable in the absence of *pknB.* However, it is clear that the absence of PknB-mediated phosphorylation of WalR causes a partial impairment of dimerization in comparison to the wild type and that this has phenotypic consequences in terms of autolysin function and antibiotic sensitivity and resistance.

We suggest that, like other response regulators, unphosphorylated WalR samples various conformations including both a closed and open conformation and, when open, a transient dimer. Phosphorylation by WalK at D53 would be expected to favor the open conformation, exposing the α4-β5-α5 interface and thereby triggering greater occupancy of the dimeric form. However, unlike all other response regulators known at this time, a secondary phosphorylation at T101 adds further stability to the dimeric form. We suggest that this occurs through the formation of a hydrogen bond between the phosphate at T101 and E108. Indeed, phosphothreonine is a relatively more stable modification than phosphoaspartate, perhaps this secondary modification serves to extend the expression of WalR target genes longer term. Replacement of T101 with a methionine eliminates phosphorylation and may substitute the important T101∼P-E108 interaction with an E108-Y99 interaction, distorting the protein and further impairing function. This would explain the greater severity of the mutant phenotype WalR_T101M_ compared to Δ*pknB*.

The evolutionary significance of this secondary modification is particularly intriguing. In addition to providing the possibility of two ‘levels’ of WalR activity, it also links transcriptional activation of WalR target genes to two biological inputs. WalK has an extracellular CACHE domain and an intracellular PAS domain. The ligand(s) for the CACHE domain are unknown but the PAS domain has been shown to bind zinc (11). Some studies have reported that PknB interacts with muropeptides, components of the cell wall, though with low binding affinity (23,43,44,48–50). PknB has also been shown to bind lipid II, a key precursor for peptidoglycan synthesis that contains all the signatures of the muropeptide ligand and is localized to the same cellular niche as PknB (23,43). Coupling the action of the WalR regulon to these critical extracellular features of cell growth likely confers a selective advantage to the cell. Future studies on the interplay between the ligands that act on WalK and PknB will likely clarify the significance of these two regulatory inputs on transcriptional regulation by WalR.

## Materials and Methods

### Strains, oligonucleotides, media, and reagents

Bacterial strains and plasmids used in this study are listed in **Table S1**. Oligonucleotides (Thermo Fisher and IDT) are listed in **Table S2**. All *S. aureus* and *E. coli* strains were grown in Tryptic Soy Broth (TSB, Oxoid) Agar or Luria Broth (LB) Agar, respectively, unless otherwise stated. When cultured in broth, they were grown in TSB or LB at 37°C with shaking at 220 rpm. Chemicals used in this study were purchased from Sigma-Aldrich unless stated otherwise. Tunicamycin was purchased from Cayman Chemical (Michigan, USA). For selection, antibiotics were added at the following concentrations: ampicillin 100 μg/mL, kanamycin 50 μg/mL, and chloramphenicol 10 μg/mL for *E. coli*; kanamycin 50 μg/mL and chloramphenicol 10-12 μg/mL for *S. aureus.* Restriction enzymes and cloning enzymes/reagents were purchased from New England Biolabs, Phusion DNA Polymerase, DreamTaq Hot Start and Phire Hot Start II DNA Polymerase were purchased from Thermo Fisher. All experiments were performed in biological triplicates with at least 3 technical replicates unless stated otherwise.

### Site-directed mutagenesis by allelic exchange of *S. aureus* mutants

To make chromosomal mutants, upstream and downstream regions for *pknB* deletion (IM255/IM1648/IM1649/IM258) were PCR amplified, next a spliced overlap extension-PCR (SOE-PCR) was performed to create an amplicon from the gel-extracted templates, which was used for SLiCE cloning into pIMAY-Z. This created plasmid *p*IMAY-Z: Δ*pknB*. Allelic exchange was performed in ATCC29213 to generate isogenic mutants as previously described (51). Putative mutants were screened by colony PCR with IM255/IM258 primers for *pknB* deletion and confirmed the mutation by whole genome sequencing. Briefly, gDNA was extracted from 1 mL of an overnight culture in BHI broth (One-Tube Bacteria Genomic DNA Isolation Kit, Bio Basic Inc.), to lyse open *S. aureus,* cultures were incubated with lysostaphin (Sigma-Aldrich, L4402) (200 μg/mL in 20 mM Tris-HCl [pH 7.5] and 10 mM EDTA). Samples were sequenced on a GridION (Oxford Nanopore Technologies) and Illumina NextSeq by the Centre for Pathogen Genomics (CPG) Innovation Hub (University of Melbourne). Resultant reads were mapped to ATCC29213 genome (GenBank accession: CP078521) using Geneious Prime 2024.0.5 (Dotmatics) and SNPs identified using Snippy (https://github.com/tseemann/snippy).

### Microtiter broth dilution assays

Cultures were grown overnight and subcultured the next day in 1:1,000 dilution in fresh media. Cultures for complementation with pRMC2 were grown in 12 μg/mL chloramphenicol and 10 ng/mL anhydrotetracycline (ATc, Cayman Chemical). Cells were grown at 37°C until exponential phase (OD600 nm of 0.4-0.5), and then subcultured in 1:10,000 dilution in fresh media. 198 μL of culture was aliquoted each into a 96-well plate and incubated with 2 μL of drug (1/100 final dilution of drug). A vehicle control and blank media control were added. Plates were incubated overnight at 37°C with shaking and read the turbidity the next day with an absorbance at 600 nm using EPOCH (BioTek) and CLARIOstar (BMG Labtech) plate readers.

### Cloning of pRMC2 constructs

*pknB* gene was PCR amplified from ATCC29213 using MS09/MS10 primers to include KpnI and EcoRI cut sites and RBS. Amplicon and pRMC2 were digested with KpnI and EcoRI, ligated using NEBuilder HiFi DNA Assembly (New England Biolabs), transformed into *E. coli* DH5α and plated on LB with ampicillin 100 μg/mL. Positive clones were screened by colony PCR and passage through *S. aureus* RN4220 and wild type ATCC29213 electrocompetent cells before electroporating into the final strain and selected for on TSB with chloramphenicol 12 μg/mL. Plasmids were miniprepped and confirmed by sequencing the insert using Sanger sequencing (TCAG, The Hospital for Sick Children, Toronto), and whole-genome sequencing using a GridION (Oxford Nanopore Technologies, CPG Innovation Hub, University of Melbourne).

### Growth curve and colony-forming unit assays

5-mL cultures were grown overnight in LB, subcultured the next day to an OD600 nm of 0.05 in fresh media into a clear sterile 96-well plate. Growth was measured every 10 min for 8 h at 37°C with dual-orbital shaking at 300 rpm in a CLARIOstar plate reader (BMG Labtech).

### *In vivo* autolysis activity assays

An overnight culture was grown and subcultured the next day in 1:1,000 dilution in 25-mL of fresh media. Cultures for complementation with pRMC2 were grown in 12 μg/mL chloramphenicol and 10 ng/mL ATc. Once cells reached early exponential phase (OD600 nm of 0.3-0.4), cells were chilled on ice immediately and harvested by centrifugation at 3,500 x g for 15 min at 4°C. Pellets were washed twice with cold sterile double-distilled water (ddH_2_O). Next, pellets were resuspended in 50 mM Tris-HCl (pH 7.4) and 0.05% (v/v) Triton X-100 to an OD600 nm of 1. Samples were aliquoted in triplicates into a clear sterile 96-well plate, and OD600 nm was measured every 30 min for 6 h at 37°C with shaking (200 rpm) in Synergy (BioTek) and CLARIOstar (BMG Labtech) plate readers.

### Overexpression and purification of proteins

For production of WalR and WalR_T101M_, full-length protein was amplified from ATCC29213 or WalR_T101M_ gDNA and cloned into pET-21a(+) plasmid with primers MS03/MS04, carrying a C-terminal 6xHis tag. For production of WalR_T101A_ and WalR_T101S_ variants, genes were synthesized and cloned into pET-21a(+) plasmids by GenScript Biotech Corporation (Nanjing, China). For production of the cytoplasmic region of PknB kinase (1-348 aa), the specific region was amplified using MS01/MS02 primers and cloned into pET-28a(+) plasmid with an N-terminal 6xHis tag. Constructs were transformed into *E. coli* BL21(DE3); cultures were grown in 1L of LB broth at 37°C with shaking until they reached OD600 nm of 0.6-0.7, and then induced with 1 mM IPTG for 4 hours at 37°C. Next, cell pellets were harvested by centrifugation. For purification of proteins, cell pellets were resuspended in binding buffer [20 mM Tris-HCl (pH 7.6), 200 mM NaCl, 5 mM imidazole, 5 mM β-mercaptoethanol] supplemented with EDTA-free protease inhibitor cocktail (1 tablet/1L culture; Roche, COEDTAF-RO). Cells were lysed by sonication, and lysates were clarified by centrifugation at 30,000 x g for 15 min at 4°C. The supernatant was separated from the cell debris, and 800 μL of Ni-NTA bead slurry (Qiagen) was added to the mixture (bead slurry had been previously equilibrated with binding buffer). Supernatants were incubated with the beads in a rotary platform for 30 min at 4°C to promote protein binding. Next, supernatants were loaded onto a 50-mL free-flow gravity column (Bio-Rad) and washed with 10 mL of wash buffer [20 mM Tris-HCl (pH 7.5), 200 mM NaCl, 30 mM imidazole, 5 mM β-mercaptoethanol]; wash step was repeated 5 times or until eluate didn’t turn blue upon addition of Bradford reagent (Bio-Rad, 1/10 dilution). Proteins were eluted with 2 mL of elution buffer [20 mM Tris-HCl (pH 7.5), 200 mM NaCl, 300 mM imidazole, 5 mM β-mercaptoethanol]; various elutions were collected separately, and all were dialyzed overnight with a 6-8 kDa MWCO dialysis bag (Spectrum Spectra/Por) in 4L of dialysis buffer [25 mM Tris-HCl (pH 7.5), 100 mM NaCl, 10% glycerol]. Protein concentration was measured using Qubit Protein Broad Range Assay Kit (Thermo Fisher), concentrated if needed using Amicon Ultra Centrifugal Filters 10 kDa MWCO (Millipore), aliquoted, snap frozen in liquid nitrogen, and stored at -80°C until required.

### Radioactive *in vitro* phosphorylation assays

Purified PknB (3 μg), alone, or with different WalR proteins (1:4 ratio, kinase: regulator) were incubated with 2.5 μCi [y-^32^P]-ATP and 5 μM ATP in 25 μl of phosphorylation buffer [25 mM Tris-HCl (pH 8), 300 mM NaCl, 0.5mM DTT, 20 mM KCl, 2 mM MgCl_2_] at room temperature. Reaction was stopped with 5X SDS-loading dye every 5 min to a maximum of 20 min. Samples were loaded in a 12% SDS-PAGE gel and run for 30 min at 180 V in 1X Tris-Glycine-SDS buffer. Next, gel was transferred to a Whatman filter paper, wrapped with plastic film and gel was exposed to a storage phosphor screen (Molecular Dynamics) overnight. Phosphor screen was imaged with a Typhoon FLA 9500 imager (GE Healthcare), and images were analyzed with ImageJ (U.S. National Institute of Health).

### *In vitro* phosphorylation assays with SEC-MALS

To characterize WalR/WalR_T101M_ oligomeric state upon T101 phosphorylation, size-exclusion chromatography coupled to multi-angle light scattering (SEC-MALS) was performed. Briefly, purified PknB (3 μg), alone, or with WalR/WalR_T101M_ proteins (1:15 ratio, kinase: regulator) were incubated with or without 500 μM ATP in phosphorylation buffer [25 mM Tris-HCl (pH 8), 300 mM NaCl, 0.5 mM TCEP, 20 mM KCl, 10 mM MgCl_2_] overnight at 4°C. Next day, the reaction was supplemented with 500 μM ATP and further incubated at 30°C for 2 h. Reactions were subjected to SEC-MALS using Advanced SEC 300 Å, 4.6 x 300 mm, 2.7 μm, SEC column (PL1580-5301, Agilent) in running buffer [25 mM Tris-HCl (pH 8), 300 mM NaCl, 0.5 mM TCEP]. Proteins were run for 35 min at room temperature. Runs were analyzed with ASTRA software (Wyatt technologies). Experiments were performed at SPARC BioCentre, Hospital for Sick Children, Toronto, Canada.

### Cloning of split luciferase vectors

Cloning of the different WalR variants and PknB (cytoplasmic domain only) in pSmBIT and pLgBIT was performed as described previously (18). Briefly, both plasmids were digested with KpnI, gel extracted and used as template for PCR with IM515/IM1360 primers. WalR alleles were PCR amplified with IM1363/IM1364 from ATCC29213, WalR_T101M_ gDNA, or pET-21a(+) plasmids; and PknB was PCR amplified with MS30/MS31 from gDNA. For WalR_E108A_-SmBIT and WalR_E108A_-LgBIT constructs, first the E108A mutation was made by SOE-PCR with primers IM1363/MS32/MS33/IM1364. For WalR_D53A_-SmBIT/WalR_D53A_-LgBIT and WalR_D53A/T101A_-SmBIT/WalR_D53A/T101A_-LgBIT constructs, first the D53A mutation was made by SOE-PCR with primers IM278/IM279 in wild type WalR to create WalR_D53A_, and in WalR_T101A_ to create WalR_D53A/T101A_. The amplicons were SLiCE cloned into pSmBIT and/or pLgBIT and transformed into IM08B with ampicillin selection. At least 1 μg of plasmid was purified from IM08B and co-electroporated into ATCC29213 wild type or the Δ*pknB* mutant with selection on BHI agar containing 10 μg/mL chloramphenicol (pSmBIT) and 50 μg/mL kanamycin (pLgBIT).

### Measurement of luciferase activity

Luciferase assays were performed as previously described (18). Briefly, 5-mL cultures in LB containing antibiotics (in a 50-mL tube) were grown overnight. The cultures were diluted 1:100 in fresh LB containing antibiotics and a 1:5,000 dilution of Nano-Glo Luciferase Assay Substrate (Promega). A 200 μL aliquot was dispensed in duplicate into a black/clear bottom 96-well plates (165305, Thermo Fisher). The plates were sealed with MicroAmp Optical Adhesive Film (Thermo Fisher) and incubated at 37°C with dual-orbital shaking at 300 rpm in a CLARIOstar plate reader (BMG Labtech). Light emission (1 s exposure) and culture growth (OD_600_) were measured every 10 min for 10 h.

### Transmission electron microscopy

A 25-mL TSB overnight culture was grown and sub-cultured the next day in 1:1,000 dilution in fresh media. Cells were grown until exponential phase (OD600 nm = ∼0.6) and harvested by centrifugation at 3,000 x g for 5 min. Samples were processed for TEM imaging as previously described (16). Briefly, samples were primary fixed with 4% paraformaldehyde and 1% glutaraldehyde solution at room temperature for 1 h followed by an overnight incubation at 4°C. Next, samples were secondary fixed with 1% osmium tetroxide, stained with 1% uranyl acetate and dehydrated using increasing concentrations of ethanol (30%, 50%, 70% and 95%). Between each step, samples were washed with ddH_2_O. Samples were washed with propylene oxide and infiltrated with Spurr’s resin and propylene mixture accordingly: 50% resin for 30 min, 75% resin for 1 h, and 100% resin overnight at room temperature with rotation. Resin-embedded samples were placed in polyethylene BEEM capsules and left to polymerize with fresh resin at 60°C for 48 h. Samples were sectioned on a Reichert Ultracut E microtome to 100 nm thickness and collected on 300 mesh copper grids. Sections were counter-stained with Reynold’s lead citrate for 10 min. Micrographs were captured using a TALOS L120C electron microscope at an accelerating voltage of 120 kV (Thermo Fisher Scientific). Images were captured at 22,000X magnification, and were processed and analyzed with ImageJ (U.S. National Institute of Health).

### CD thermal melt assay

To assess changes in protein stability of WalR_T101M_ in comparison to wild type WalR, thermal melt assays were performed using circular dichroism (CD). Briefly, purified proteins were diluted in running buffer [25 mM Tris-HCl (pH 7.5), 100 mM NaCl] to achieve concentration of 0.2 mg/mL, and single-wavelength measurements were taken for a temperature range of 20-95°C at 222 nm (this specific wavelength was chosen based on a preliminary CD spectrum analysis of the proteins that produced mostly an α-helix shape). A blank with only running buffer was also included to extract background noise from the samples. Experiments were performed in a Jasco J-1500 Spectropolarimeter (Jasco Inc), and only one technical replicate was run for each protein.

### Modeling of WalR receiver domain of *S. aureus*

The coding sequence corresponding to the WalR receiver domain (residues 1-122; UniProtKB entry Q7A8E1) was used for three-dimensional structure prediction. Homology modeling was performed using the Swiss-Model web server (47) to generate an initial structural model. The crystal structure of WalR receiver domain from *Bacillus subtilis* (PDB: 2ZWM) was selected as the template because it provides explicit structural information on the dimer interface. Post-modeling refinement was conducted using Schrödinger Maestro version 2021 (Schrödinger, Inc., USA) in combination with the “BioLuminate” module. The refinement process involved energy minimization using the OPLS3e force field (52). Specific post-translational modifications were introduced to represent the functional state of the dimer: aspartate 53 was phosphorylated on both chains, as well as the phosphothreonine at residue T101. These modifications were implemented using the “Build” module in Schrödinger Maestro while maintaining structural integrity. The quality of the homology model was assessed using the “Structure Assessment” feature of the Swiss-Model platform (53), with tools such as Ramachandran plot analysis, MolProbity analysis (54), and QMEANDisCo scoring (55–57), ensuring that the predicted model was suitable for subsequent interaction analyses. Briefly, the Ramachandran plot revealed that 97.35% of the residues were located in favored regions, with no residues present in outlier regions. MolProbity score of the model was 0.93, reflecting high structural quality and reliability and a low clash score of 1.04. The absence of rotamer outliers and C-beta deviations further supports the structural integrity of the model. The QMEANDisCo global score of the model was 0.81 ± 0.05, falling within the acceptable range for high-quality homology models. All of these analyses suggested that the model was an accurate depiction of WalR dimerisation and suitable for further analysis. The finalized structure served as the wild type reference for all downstream computational experiments.

### Interaction analysis of dimerization interface of WalR

To evaluate the importance of T101 on the dimerization interface, and the impact of T101M and T101A mutations on the overall dimer stability, we created these two additional variants. These mutations were introduced using the “Build” module in Schrödinger Maestro, maintaining the phosphorylated state of D53 in both structures. The three final dimers—wild type T101∼P, T101M, and T101A—underwent structural refinement using the “Protein Preparation Wizard” in Schrödinger Maestro. This preparation included the addition of hydrogen atoms, and optimization of hydrogen bonding networks. The OPLS3e force field was then employed to minimize the energy of each model and eliminate any steric clashes or unfavorable contacts (52). Next, “Protein Interaction Analysis” tool in BioLuminate was used to examine intermolecular interactions in detail. We analyzed any hydrogen bonds, salt bridges, and hydrophobic contacts involving T101 residue and its neighboring residues. The same was conducted in the models for T101M and T101A.

### Molecular dynamics simulations

The three dimer structures were subjected to molecular dynamics (MD) simulations using the “Desmond” module in Schrödinger Maestro. Each dimer structure was individually placed in an orthorhombic simulation box, which was then filled with explicit TIP3P water molecules (58). Counter ions were introduced to neutralize the net charge of the system. Prior to initiating the production run, the system underwent an initial energy minimization step to remove any unfavorable contacts, followed by a heating phase that gradually raised the temperature from a lower value to 300 K. This was succeeded by an equilibration phase under constant temperature (300 K) and pressure (1 atm) conditions, maintained by the Nose–Hoover thermostat and barostat (59). The simulations were particularly focused on capturing the hydrogen bonds at residue 101 within the dimer interface. Upon completion of the production MD runs, the trajectory was analyzed to identify how residue T101 influences intermolecular interactions and overall dimer stability across the wild type and mutant structures. Videos illustrating these interactions for each WalR dimer are provided in the Supplementary Data (**Movie 1-3**).

### Graphical and statistical analysis

Statistical analyses were performed using GraphPad Prism software, version 9.2.0 (GraphPad Software, LLC). Statistical significances were calculated using one-way ANOVA when comparing more than two groups, and non-parametric Wilcoxon matched-pairs signed rank test and/or Paired t test (parametric values) were used to compare two groups only. Area under curve was calculated to compare peak luminescence values between samples. All values in the graphs are expressed as the median and error (minimum to maximum values) unless otherwise stated.

## Data availability

All data are available within the article and supporting information. All material and correspondence should be directed to and/or.

## Acknowledgments

This work was supported by Canadian Institutes for Health Research grant #202209 from to J.R.N. and by Australian National Health and Medical Research Council Ideas grant #GNT2021638 to S.J.P., and Australian Research Council Discovery Project Grant #DP230102668 to T.P.S. and S.J.P.; M.S. was supported by a University of Melbourne/University of Toronto joint PhD scholarship. M.S. is a fellow in the joint University of Toronto/University of Melbourne PhD program.

## Author contributions

Conceptualization, M. S., J. R. N., S. J. P., I. R. M., L. K. R. S., T. P. S.; Methodology, M. S., I. R. M., L. K. R. S., N. E. S., D. G., A. N., S. T., K. M., S. M. P.-E.; Investigation, M. S., I. R. M., N. E. S., D. G., A. N.; Formal Analysis, M. S., S. J. P., L. K. R. S., D. G.; Writing-Original Draft, M. S.; Writing-Review and Editing, M. S., J. R. N., S. J. P., I. R. M., L. K. R. S., T. P. S.; Visualization, M. S.; Funding Acquisition, J. R. N. and S. J. P.

