## Supplemental figures for "Phosphorylation by the PknB serine/threonine kinase stimulates dimer formation by WalR"

### Figure S1

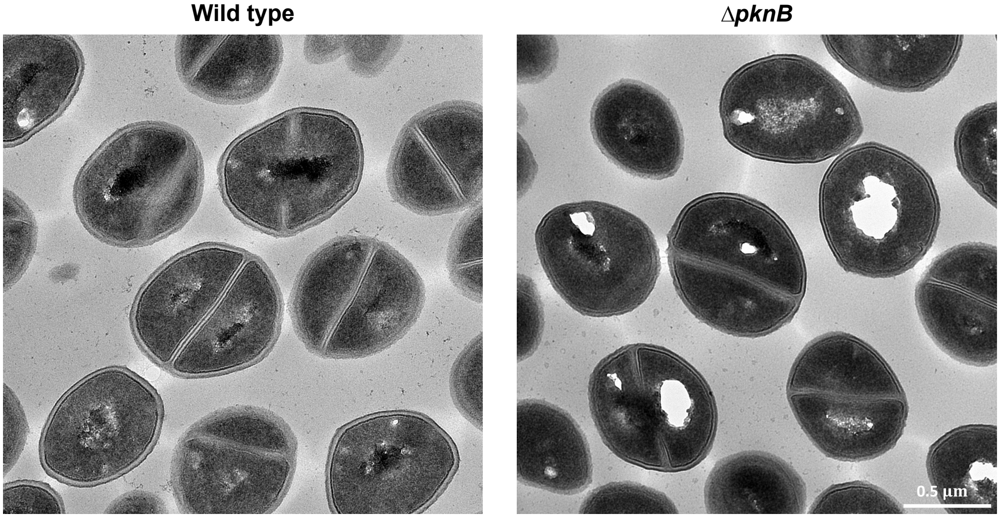


**Transmission electron micrographs of wild type and Δ*pknB* strains during log-phase growth.**

### Figure S2

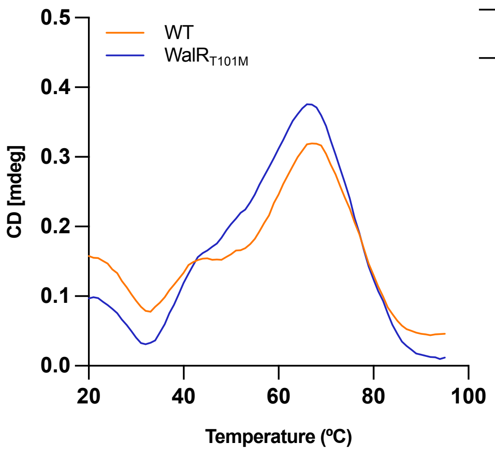


| | Melting temperature Tm (ºC) |
|---|---|
| **WT** | 67 |
| **WalR$_{T101M}$** | 66 |

**Protein stability of native WalR and WalR$_{T101M}$ proteins.** Thermal melt assays were performed to assess changes in protein stability of WalR$_{T101M}$ compared to wild type WalR, using circular dichroism (CD). Proteins were loaded at final concentration of 0.2 mg/mL, and single-wavelength measurements were taken for a temperature range of 20-95°C at 222 nm. A blank with only running buffer was also included to extract background noise from the samples. Only one technical replicate was run for each protein. Melting temperatures of both proteins are also indicated.

### Figure S3

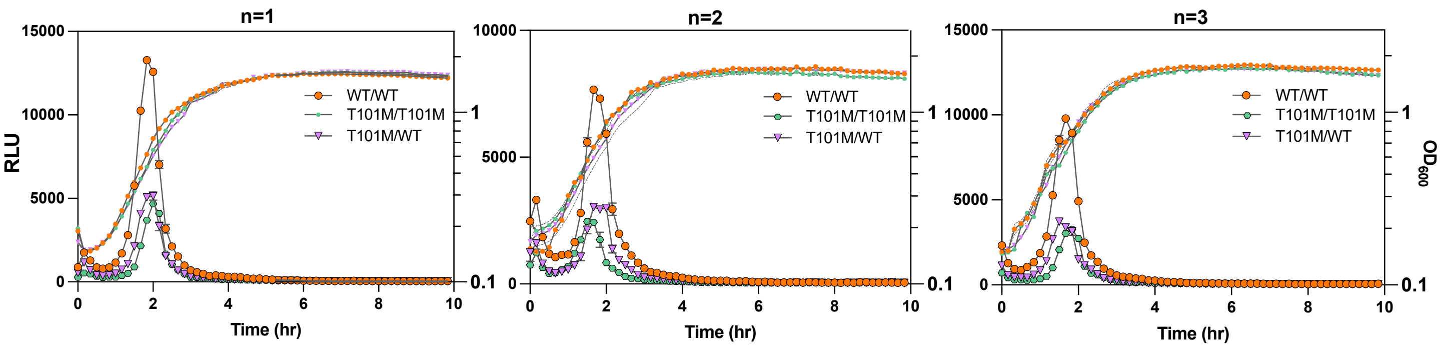


**Wild type WalR-SmBIT and WalR$_{T101M}$-LgBIT kinetic interactions.** T101M mutation was observed to be dominant and only one allele was sufficient to alter native WalR/WalR interaction. Three independent experiments were done in wild type *S. aureus*.

### Figure S4

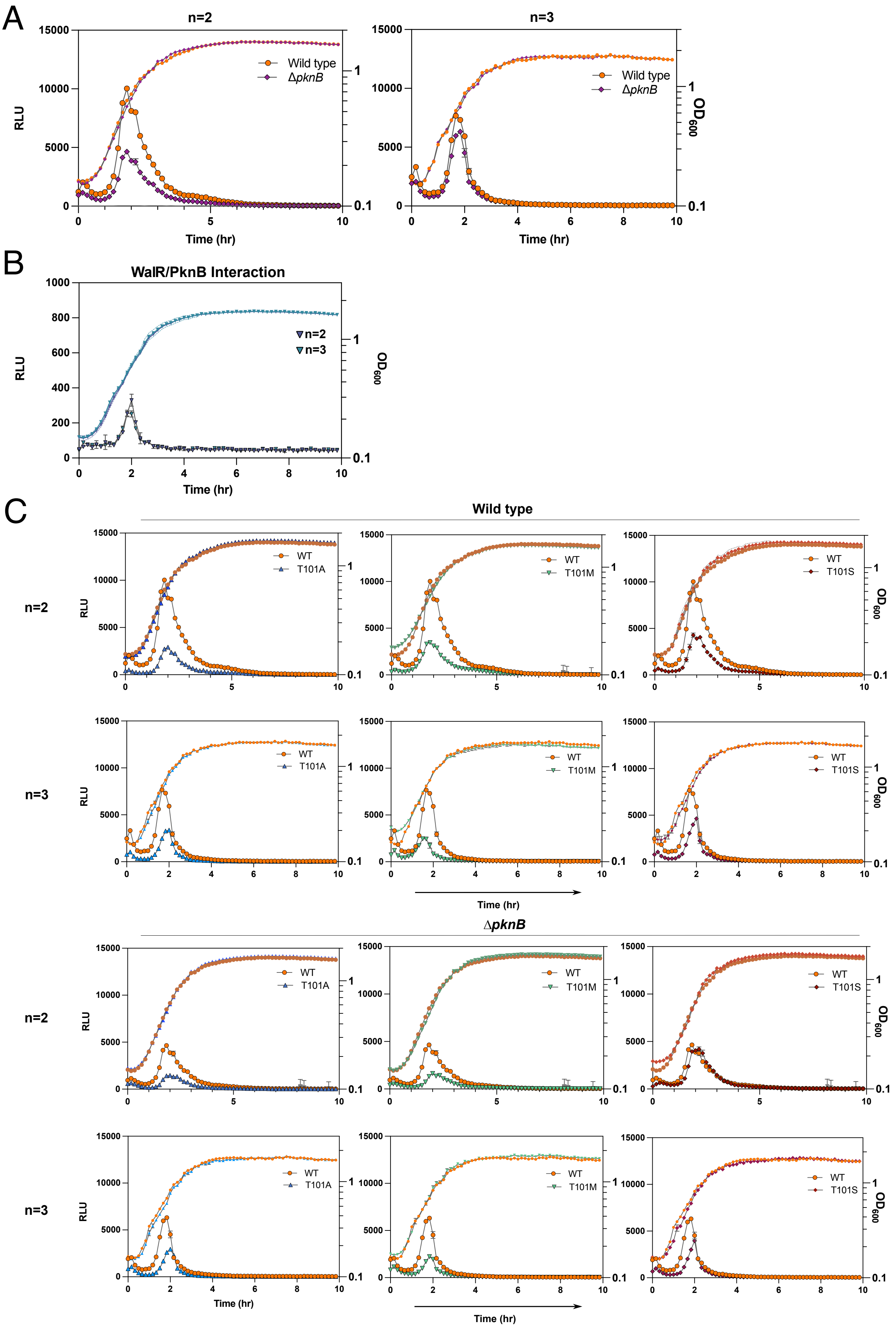


**Biological replicates of the experiments shown in Figure 3 (not shown in the main text). A)** Analysis of the interaction of WalR-SmBIT/LgBIT in wild type and Δ*pknB* backgrounds. Showing n=2 and 3. **B)** Kinetic interactions of WalR/PknB throughout growth in wild type cells. Showing n= 2 and 3. **C)** Comparison of wild type WalR/WalR kinetics to three mutant versions of WalR ($WalR_{T101M}$, $WalR_{T101A}$, and $WalR_{T101S}$) in wild type and Δ*pknB* backgrounds. Showing n= 2 and 3.

### Figure S5

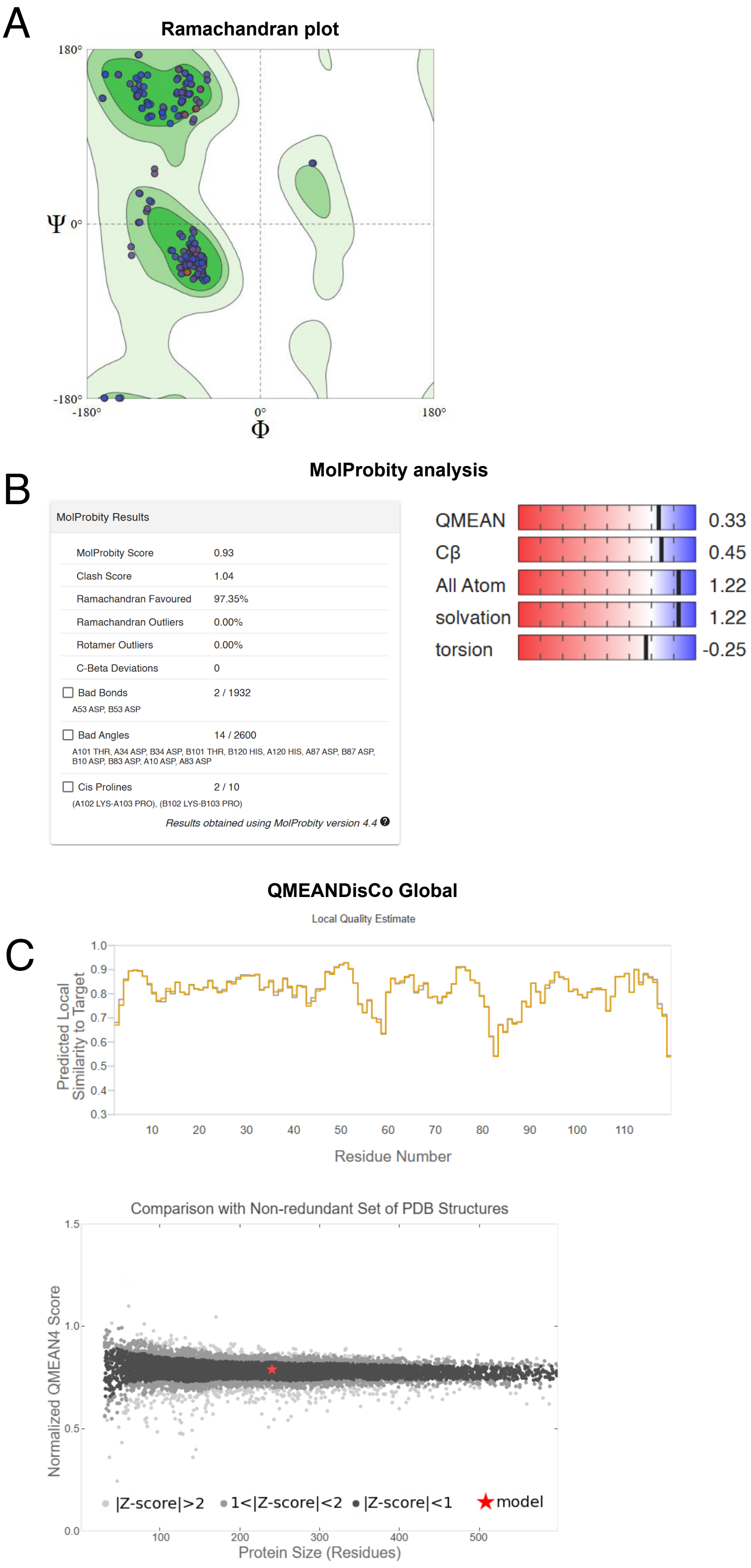


**Structural assessment of the predicted dimer of the WalR receiver domain in *S. aureus.*** Various bioinformatic tools were used to evaluate the reliability of the predicted dimer for subsequent interaction analyses. **A)** Ramachandran plot analysis. **B)** MolProbity analysis. **C)** QMEANDisCo Global scoring.

### Figure S6

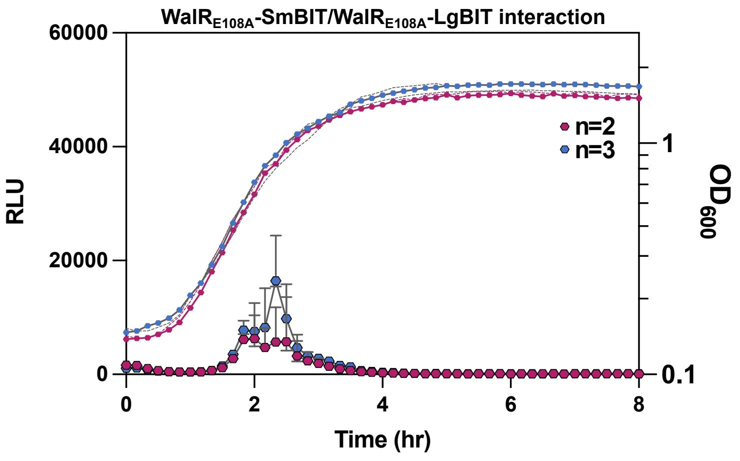


**Biological replicates of the experiments shown in Figure 4E (not shown in the main text).** Analysis of the interaction of WalR$_{E108A}$-SmBIT/WalR$_{E108A}$-LgBIT in Δ*pknB* background. Showing n=2 and 3.

### Figure S7

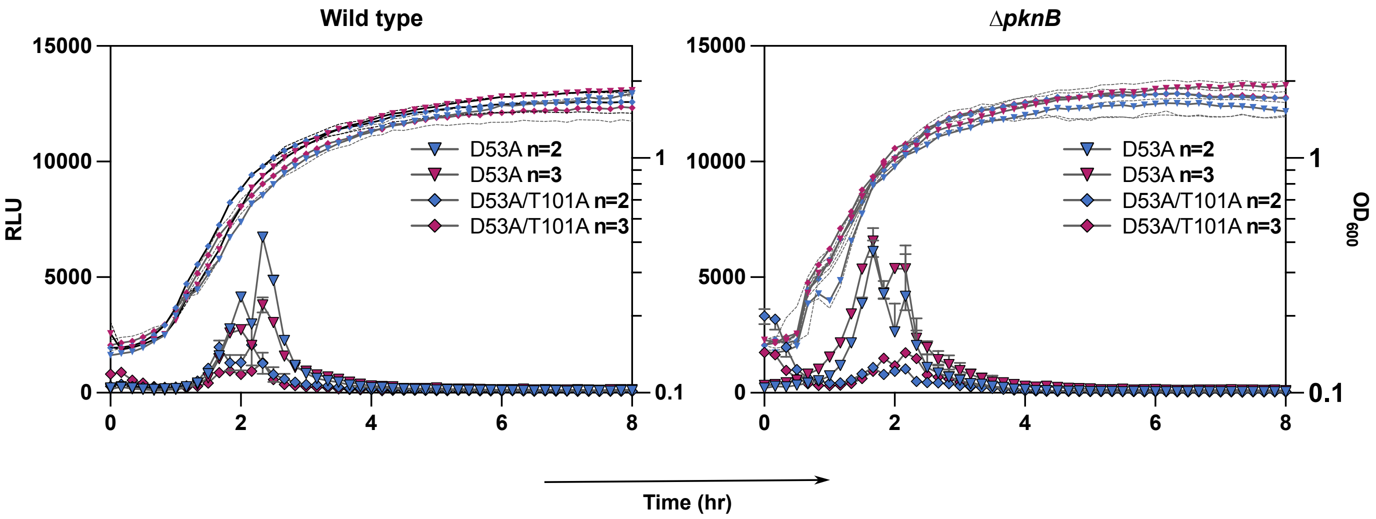


**Biological replicates of the experiments shown in Figure 5 (not shown in the main text).** Analysis of the interaction of WalR$_{D53A}$-SmBIT/WalR$_{D53A}$-LgBIT and WalR$_{D53A/T101A}$-SmBIT/WalR$_{D53A/T101A}$-LgBIT in wild type and Δ*pknB* backgrounds. Showing n=2 and 3.

### Table S1

| ***Staphylococcus aureus* strains** | **Description** | **Source** |
|---|---|---|
| ATCC 29213 | *Staphylococcus aureus* | ATCC |
| RN4220 | Restriction deficient *S. aureus* cloning strain; mutation in sau1 and hsdR; mec-, rsbU- and agr- | Lab stock |
| $WalR_{T101M}$ | *S. aureus* ATCC29213 with $WalR^{T101M}$ chromosome mutation. Alternatively called *walR1.* | Lab stock |
| Δ*pknB* | *S. aureus* ATCC29213 with clean deletion of *pknB* gene from ATG to TAA. | This study |
| ***Escherichia coli* strains** | **Description** | **Source** |
| *E. coli* DH5**α** | K12 stain. *fhu*A2 *recA1 endA1 hsdR17.* Chemically competent strain for high efficiency transformation, and cloning. | New England Biolabs |
| *E. coli* BL21(DE3) | B strain. *fhuA2 [lon] ompT gal (λ DE3) [dcm] ∆hsdS.* IPTG-inducible T7 RNA polymerase. Chemically competent strain for protein production. | New England Biolabs |
| IM08B | DH10BΔ*dcm.* Expresses CC8 adenine methylation profile for direct transformation of ATCC 29213. | (1) |
| **Plasmids** | **Description** | **Source** |
| pET-21a (+) | IPTG inducible protein expression plasmid. Ampicillin resistant. | Lab stock (Addgene) |
| pET-28a (+) | IPTG inducible protein expression plasmid. Kanamycin resistant. | Lab stock (Addgene) |
| pRMC2 | Anhydro-tetracycline inducible plasmid for *S. aureus* manipulation. | (2) |
| pIMAY-Z | Allelic exchange plasmid. Cm(R) | (1) |
| pSmBIT | pRAB11 backbone (pC194 replicon) with SmBIT split luciferase. Ampicillin(R) Chloramphenicol(R). | (3) |
| pLgBIT | pCN34 backbone (pT181 replicon). TetR/tetO ex pRAB11 introduced. LgBIT split luciferase. Ampicillin(R) Kanamycin(R). | (3) |
| **Protein overexpression in *E. coli*** | **Description** | **Source** |
| pET21a: *walR* | For production of recombinant WalR in *E. coli.* C-terminally tagged with 6xHis tag. | This study |
| pET21a: $walR_{T101M}$ | For production of recombinant $WalR_{T101M}$ in *E. coli.* . C-terminally tagged with 6xHis tag. | This study |
| pET21a: $walR_{T101A}$ | For production of recombinant $WalR_{T101A}$ in *E. coli.* . C-terminally tagged with 6xHis tag. | GenScript |
| pET21a: $walR_{T101S}$ | For production of recombinant $WalR_{T101S}$ in *E. coli.* . C-terminally tagged with 6xHis tag. | GenScript |
| pET28a: $pknB_{CYTO}$ | For production of recombinant cytoplasmic region of PknB (1→348 aa) in *E. coli.* N-terminally tagged with 6xHis tag, T7 tag and thrombin site. | This study |
| **Gene complementation plasmids for *S. aureus*** | **Description** | **Source** |
| pRMC2: *pknB* | ATc inducible plasmid containing of *pknB* gene from ATCC 29213, for complementation. | This study |

| **Split luciferase plasmids for protein: protein interaction in *S. aureus*** | **Description** | **Source** |
|---|---|---|
| pSmBIT: *walR* | C-terminally tagged WalR with SmBIT luciferase fragment. | This study |
| pSmBIT: $walR_{T101M}$ | C-terminally tagged $WalR_{T101M}$ with SmBIT luciferase fragment. | This study |
| pSmBIT: $walR_{T101A}$ | C-terminally tagged $WalR_{T101A}$ with SmBIT luciferase fragment. | This study |
| pSmBIT: $walR_{T101S}$ | C-terminally tagged $WalR_{T101S}$ with SmBIT luciferase fragment. | This study |
| pSmBIT: $walR_{E108A}$ | C-terminally tagged $WalR_{E108A}$ with SmBIT luciferase fragment. | This study |
| pSmBIT: $walR_{D53A}$ | C-terminally tagged $WalR_{D53A}$ with SmBIT luciferase fragment. | This study |
| pSmBIT: $walR_{D53A/T101A}$ | C-terminally tagged $WalR_{D53A/T101A}$ with SmBIT luciferase fragment. | This study |
| pLgBIT: *walR* | C-terminally tagged WalR with LgBIT luciferase fragment. | This study |
| pLgBIT: $walR_{T101M}$ | C-terminally tagged $WalR_{T101M}$ with LgBIT luciferase fragment. | This study |
| pLgBIT: $walR_{T101A}$ | C-terminally tagged $WalR_{T101A}$ with LgBIT luciferase fragment. | This study |
| pLgBIT: $walR_{T101S}$ | C-terminally tagged $WalR_{T101S}$ with LgBIT luciferase fragment. | This study |
| pLgBIT: $walR_{E108A}$ | C-terminally tagged $WalR_{E108A}$ with LgBIT luciferase fragment. | This study |
| pLgBIT: $pknB_{CYTO}$ | C-terminally tagged PknB (cytoplasmic domain only, 1→348 aa) with LgBIT luciferase fragment. | This study |
| pLgBIT: $walR_{D53A}$ | C-terminally tagged $WalR_{D53A}$ with LgBIT luciferase fragment. | This study |
| pLgBIT: $walR_{D53A/T101A}$ | C-terminally tagged $WalR_{D53A/T101A}$ with LgBIT luciferase fragment. | This study |
| **Allelic exchange constructs for *S. aureus*** | **Description** | **Source** |
| pIMAY-Z: *pknB* | For deletion of *pknB* gene by allelic exchange. | This study |

### Table S2

| Primer | Sequence (5'→3') | Name |
|---|---|---|
| MS01 | cagagcaaatgggtcgcGGATCCataggtaaaataataaatgaacg | *pknB* gene fwd with BamHI site. Cloning and protein overexpression. |
| MS02 | tggtggtggtgtcaCTCGAGctttcgtgttgatttcttttagg | *pknB* gene rev with XhoI site. Cloning and protein overexpression. |
| MS03 | gaaggagatataCATATGgctagaaaagttgttgta | *walR* gene fwd with NdeI site. Cloning and protein overexpression. |
| MS04 | gtgggtggtggtgCTCGAGctcatgttgttgga | *walR* gene rev with XhoI site. Cloning and protein overexpression. |
| MS05 | gctaaagctttaagtgagacgtctttaactcaga | *pknB* screening fwd. Colony PCR-screening of insert. |
| MS06 | agtttctccattaaagggtggttcaccaacaagc | *pknB* screening rev. Colony PCR-screening of insert. |
| MS07 | cagcagccaactcagttcct | *walR* screening fwd. Colony PCR-screening of insert. |
| MS08 | tcgagatctcgatcccgcg | *walR* screening rev. Colony PCR-screening of insert. |
| MS09 | attaaaataagcttgatGGTACCaggaggtatatacatatgataggtaaaataataaatgaac | *pknB* gene fwd with KpnI and RBS site. For complementation purposes. |
| MS10 | tgtaaaacgacggccagtgaattcttaaatatcatcatagctgacttc | *pknB* gene rev with EcoRI site. For complementation purposes. |
| MS11 | atcccctcgagttcatgaa | pRMC2 screening fwd. Colony PCR-screening. |
| MS12 | atactcatgtgctgcaaggc | pRMC2 screening rev. Colony PCR-screening. |
| MS20 | aatacctgtgacggaagatcacttcg | pIMAY-Z screening fwd. Colony PCR-screening. |
| MS21 | tacatgtcaagaataaactgccaaagc | pIMAY-Z screening rev. Colony PCR-screening. |
| MS22 | ggtacccagcttttgttccctttagtgagg | pIMAY-Z plasmid amplification fwd. |
| MS23 | gagctccaattcgccctatagtgagtcg | pIMAY-Z plasmid amplification rev. |
| IM1363 | gatagagtatgatgaggaggaattggaaaatggctagaaaagttgttgtagttg | *walR* gene pSm/LgBIT fwd. Cloning into pSm/LgBIT plasmids. |
| IM1364 | gaaccaccaccaccactagaaccctcatgttgttggaggaaatatcc | *walR* gene pSm/LgBIT rev. Cloning into pSm/LgBIT plasmids. |
| IM515 | tttccaattcctcctcatcatactctatc | pSm/LgBIT plasmid amplification rev. |
| IM1360 | ggttctagtggtggtggtggttctgg | pSm/LgBIT plasmid amplification fwd. |
| IM255 | cctcactaaagggaacaaaagctgggtaccatgagttgaaatcccgttttgaagc | Delta *pknB* A fwd primer. For allelic exchange constructs. |
| IM1648 | **tttacctatcat**actttatcaccttcaatagc | Delta *pknB* B rev primer. For allelic exchange constructs. |
| IM1649 | tattgaaggtgataaagtatgataggtaaataaatataattgaagtaaatgtaccgagg | Delta *pknB* C fwd primer. For allelic exchange constructs. |
| IM258 | cgactcactatagggcgaattggagctctcgcatttaactgatacgaatgtgc | Delta *pknB* D rev primer. For allelic exchange constructs. |

**Uppercase:** restriction enzyme sites.
**Underlined:** tails complementary to plasmid for cloning.
**Bold:** mutations introduced to make chromosomal mutants.

| Primer | Sequence (5'→3') | Name |
|---|---|---|
| MS30 | gatagagtatgatgaggaggaattggaaaatgataggtaaaataataaatgaacg | *pknB* gene pSm/LgBIT fwd. Cloning into pSm/LgBIT plasmids. Only cytoplasmic domain of PknB. |
| MS31 | gaaccaccaccaccactagaaccctttcgtgttgatttctttttagg | *pknB* gene pSm/LgBIT rev. Cloning into pSm/LgBIT plasmids. Only cytoplasmic domain of PknB. |
| MS32 | t**g**c**t**c**t**cgtactaaacggtttcgttaca | *walR*$_{E108A}$ B rev primer. Generate *walR*$_{E108A}$ to clone into pSm/LgBIT plasmids. |
| MS33 | tatgtaacgaaaccgtttagtacg**a**g**a**g**c**attaatcgcacgtgtgaaagcgaactta | *walR*$_{E108A}$ C fwd primer. Generate *walR*$_{E108A}$ to clone into pSm/LgBIT plasmids. |
| MS34 | **tg**cta**a**taatacgatgtctggttcttcttca | *walR*$_{D53A}$ and *walR*$_{D53A/T101A}$ B rev primer. Generate *walR*$_{D53A}$ and *walR*$_{D53A/T101A}$ to clone into pSm/LgBIT plasmids. |
| MS35 | aaccagacatcgtatta**t**tag**ca**atcatgttacctggtcgtgatggtatg | *walR*$_{D53A}$ and *walR*$_{D53A/T101A}$ C fwd primer. Generate *walR*$_{D53A}$ and *walR*$_{D53A/T101A}$ to clone into pSm/LgBIT plasmids. |

**Uppercase:** restriction enzyme sites.
**Underlined:** tails complementary to plasmid for cloning.
**Bold:** mutations introduced to make chromosomal mutants.

### Table S3

| Sample | Size (aa) | Condition | State | Theoretical mass (kDa) | Mass SEC-MALS (kDa) |
|---|---|---|---|---|---|
| $PknB_{CYTO}$ | 348 | + ATP | | 42.67 | 50.6 |
| WalR | 233 | - ATP | unphosphorylated | 27.8 | 31.7 |
| | 233 | + ATP | phosphorylated/ monomer | 27.8 | 31.9 |
| | | | phosphorylated/ dimer | 55.6 | 35.8 |
| WalR1 | 233 | - ATP | unphosphorylated | 27.8 | 29.9 |
| | 233 | + ATP | phosphorylated/ monomer | 27.8 | 29.3 |
| | | | phosphorylated/ dimer | 55.6 | 35.4 |
